# MooViE – Engine for single-view visual analysis of multivariate data

**DOI:** 10.1101/2024.04.26.591357

**Authors:** Anton Stratmann, Martin Beyß, Johann F. Jadebeck, Katharina Nöh

## Abstract

**Summary:** Understanding input-output relationships within multivariate datasets is an ubiquitous task in the life and data sciences. For this, visual analysis is indispensable for providing expressive summaries and preparing decision-making. We present the visual analysis approach and software `MooViE`, which is designed to strike the balance between being tailored to the specific data semantic and while being broadly applicable. `MooViE` supports the data exploration process for extracting important information from the data and captures the result in a fresh single-view visualization. `MooViE` is implemented in C++ to facilitate fast access and effective interaction with comprehensive multivariate datasets. We showcase the engine for various application fields, relevant to the life sciences.

**Availability and Implementation:** The source code is available under MIT license at https://jugit.fz-juelich.de/IBG-1/ModSim/MooViE and https://github.com/JuBiotechMooViE, with detailed documentation and usage instructions (https://moovie.readthedocs.io), as well as zenodo-archived releases (https://doi.org/10.5281/zenodo.10997388). Platform independent Docker images are also available (jugit-registry.fz-juelich.de/ibg-1/modsim/moovie/moovie).

**Contact:** Katharina Nöh

## 1 Introduction

Visualization plays a key role in modern data analysis, where “visuals” facilitate the process of uncovering relationships hidden in complex datasets and, thereby, support comprehension, knowledge mining, and decision-making (Kehrer and Hauser, 2013). Effective visual analysis tools allow data exploration through interaction, e.g., to organize, rescale, or filter data attributes, while the resulting visualization artifacts preserve the semantics underlying the datasets. For multivariate data, i.e., data with interdependent variables, comprehensive visual analytics solutions have been developed, each tailored to represent the specific semantic that is prevalent in the particular application domain (Thorvaldsdóttir et al., 2013; Oveland et al., 2015; Nusrat et al., 2019).

We here consider the common case of multivariate data with variables that are subdivided into *input* and *output* variables, a common type of data that occurs in many domains. Typical examples are screens of biological systems under various reaction conditions (Baraibar et al., 2014), assessing the influence of model parameters on an epidemic now- or forecasts (Kühn et al., 2021), determining suitable hyperparameter combinations with respect to algorithm performance metrics (Emmerich and Deutz, 2018), or evaluating the effectiveness of experimental design choices according to multiple conflicting goals (Nöh et al., 2018; Beyß et al., 2021).

Formally, datasets with input-output semantic entail a mapping, where some (given or unknown) function maps a set of *N* numeric input variables (*parameters, sources*) to a set of *M* numeric output variables (*responses, targets*), thereby instantiating relations between and among the two types of variables. Given an input-output dataset, the goal of visual analytics is to facilitate understanding of these relationships, both qualitatively and quantitatively.

To mine insights from multivariate data, several forms of information visualizations are established (Liu et al., 2014). Among the most versatile ones are scatterplot and parallel coordinates diagrams (Johansson and Forsell, 2016). Scatterplots break multidimensional data down into an array of pairwise views, which disclose correlations between two variables. However, the efficiency of this visual deteriorates with increasing variable count (Pak Chung Wong and Bergeron, 1997). Parallel coordinates, on the other hand, perform well for higher dimensional data, but are not designed to preserve input-output data semantics, thus lacking relevant information of the variables’ interrelations. Relationships between and within data variables can be visualized using chord diagrams (Gutwin et al., 2023). Herein, a data item is represented by a set of ribbons (chords), connecting one set of variables to some other set of variables, where the thickness of the ribbons encodes the strength of the relationship. Despite visually appealing, chord diagrams do not scale well with the number of data items (Holten, 2006) and, more importantly for our context, do not conserve the relations between many inputs and many outputs.

We here present the visualization approach and engine `MooViE` supporting the visual analysis of multivariate input-output data (Figure 1). Borrowing on circular diagram layout, inspired by chord diagrams popularized by the celebrated Circos software for the visual analysis of genome data (Krzywinski et al., 2009), `MooViE` scenes fuse familiar visuals, namely histograms, chord diagrams, and parallel coordinates, to overcome the shortcomings of the single approaches in representing quantitative input-output data in a single view. With various exemplary cases at hand, we show how important information from challenging domain-specific datasets is extracted with `MooViE`. Here, the expressive visualization approach delivers comprehensive visual summaries for multivariate data. A graphical user interface (GUI) facilitates interactive data exploration workflows, while a command line interface (CLI) makes embedding `MooViE` in automated analysis workflows simple. With its fast C++ core, `MooViE` is designed for handling comprehensive datasets, while offering production-ready visuals.

**Fig. 1.**
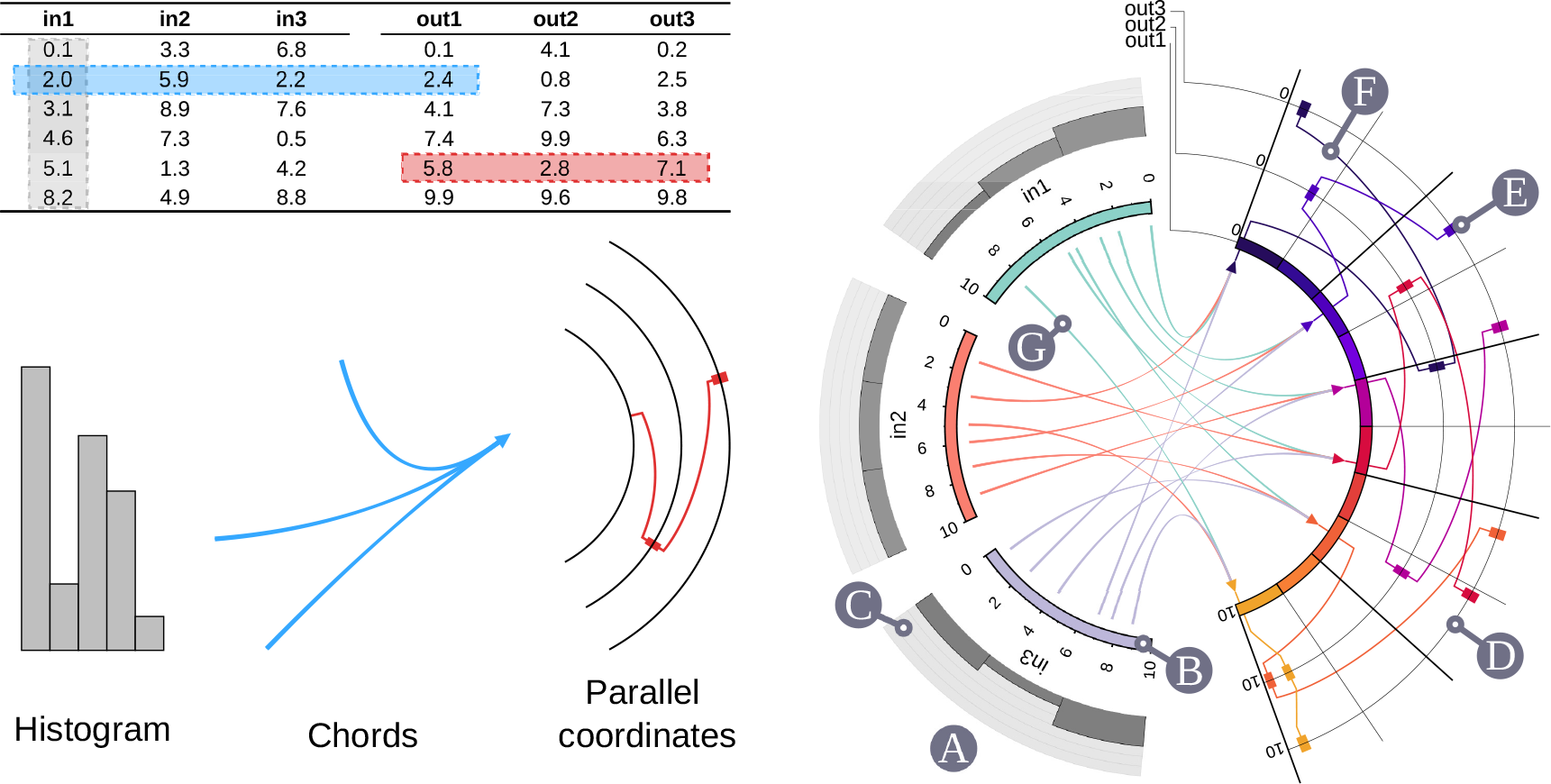
Visual anatomy and nomenclature of an example `MooViE` scene. According to the `MooViE` data model, variables are partitioned into two classes, *input* and *output* variables. Each row of the data table makes up a set of numeric values defining a *data item*. `MooViE` scenes are compiled by three forms of information visualization: The distribution of each input variable is represented by a histogram, the relationships of the input variables with the first output variable in the data table is visualized by bundled chords, and the relationship between the output variables of a data item is visualized with parallel coordinates. The nomenclature of the scene elements is: **A** input segment, **B** input arc, **C** histogram, **D** output arc where the innermost arc, assigned by out1 in the data table, is denoted primary arc, **E** box, **F** polyline, and **G** chord

## 2 Approach and Implementation

The `MooViE` engine implements a novel single-view visualization approach for multivariate data analysis, which is designed to be both fast and adaptable.

### 2.1 Data model

`MooViE`’s visualization approach is centered around the concept of data with input-output semantic. Consequently, datasets are provided as data tables, where the *input variables* populate the leading columns, followed by *output variables*. Each row of the table represents a *data item*, which defines the quantitative relationship instance between input and output variables. Formally, a dataset is a *n* × (*M* + *N*) matrix, with *n* ≥ 1 data items, *M* ≥ 1 input and *N* ≥ 1 output variables. The entries of the data table are allowed to be real- and/or integer-typed.

### 2.2 Visualization

The visuals produced by `MooViE` are circular and referred to as *scenes*. They have semantically separated left and right halves (Figure 1), being associated with input and output variables, respectively. The chords (center) and polylines (right) represent the quantitative relationship between the values of the input and output variables.

On the left of a `MooViE` scene, each input variable is assigned an individual sector, called *input segment*. Input segments consist of an *input arc*, being assigned a variable-specific color, name, and range (possibly with unit). Optionally, each segment is equipped with a *histogram* with configurable bin size, showing the (marginal) value distribution of the associated input variable. Arc colors are picked from a qualitative scale with as many classes as possible (12) using ColorBrewer (Harrower and Brewer, 2003).

On the right side, output variables are arranged in a single sector consisting of concentric arcs (*output arcs*), where each output variable resides on a single arc. Like the input segments, output arcs have a name and a range. The output arc that is located closest to the center is called *primary output arc*.

This arc acts an “attractor”, conflating the input values and connecting them to the output values of the data items. Different to the visual design of the input arcs, the primary output arc is split into 20 colored sections, ranging from dark blue (low values), over purple and red (medium), to yellow (high). Deliberately, the luminance of the colors is designed to increase, providing visual guidance for the ordering of the corresponding data values. The remaining output arcs are represented by lines. For all but the primary output variable, *boxes* are placed on the output arcs according to their values within the data item. Neighboring boxes are connected by a *polyline*. The polyline and boxes share the same color, which is determined by the value of the primary output variable. Thereby, following polylines from the primary output arc is straightforward, even when polyline cross.

Beyond the ribbons in traditional chord diagrams, the *chords* in `MooViE` scenes connect the set of input values and the value of the primary output variable, thereby representing the quantitative relation between them: Chords are connecting curves (bundles), drawn from each input segment to the primary output arc, where start and end-points of the chord bundle are given by the values of the associated input and primary output variables in a data item, respectively. Hence, all chords that belong to one data item meet at the same point of the primary output arc, where an arrowhead is placed. The chords of each data item share the same color with the input variable, while the color of the arrowhead is set to the color of the value on the primary output arc.

### 2.3 Software

Given a valid dataset, `MooViE` scenes are rendered using the visualization engine `MooViE`. The engine consists of a desktop application with graphical user interface (GUI), a command line interface (CLI) and a library.

The desktop application is designed to support interactive creation of custom visualizations and data exploration. Features include the reordering and disabling of variables, which is useful for disentangling polylines connecting output variables and resizing the ranges of individual variables, and which allows focussing on parts of the data. Further options to customize the visual appearance of `MooViE` scenes comprise, for instance, the selection of radii for input and output arcs, line thickness, fonts, and the size of the produced image. A full list of features is found in the software’s user manual available at https://moovie.readthedocs.io.

Complementing the interactive user-centric GUI, `MooViE` provides a CLI, which allows efficient generation of `MooViE` scenes where large amounts of similar datasets are to be processed, e.g. in automated data analysis pipelines. With its well-designed CLI, `MooViE` integrates seamlessly into automation scripts, where the interface offers the same options for customizing visual appearance as the GUI.

The `MooViE` library is the backend of both, the GUI and CLI, and integrates into any C++ code to provide its full functionality for data modification and visual configuration. Both GUI and library are capable of serializing the in-memory version of data tables and scene configuration as human- and computer-readable text files. Given the data table and the configuration file, `MooViE` scenes are uniquely defined and, thereby, reproducible. In particular, the configuration enables reproducibility of the results of interactive data explorations through the CLI. With that, `MooViE` supports all scenarios, from the one-off visual exploration of complex datasets to analysis pipelines at scale.

### 2.4 Implementation

The `MooViE` engine is implemented in C++ to rapidly generate images even for large data tables, where parallelized processing over multiple datasets is possible. Handling of data tables with up to *n* = 20, 000 entries is possible without performance drop (SI Text Sec. S.2). For efficient rendering, the vector graphics library Cairo is used (version 1.16, https://cairographics.org).

Input and output of `MooViE` are file-based. Data files require a tabular CSV-based format, where input and output variable columns and names are specified in the table header. To maintain visual expressiveness, `MooViE` implements upper bounds for the number of inputs (12) and outputs (8). Generated output files are vector graphics files that follow the efficient SVG Tiny 1.2 standard (W3C, 2008). Configuration files are text files consisting of lines with a key-value syntax: `moovie.<setting> = <value>`.

The GUI of the `MooViE` desktop application is written in QT (version 5 or 6, https://qt-project.org). Variable re-ordering is implemented as drag-and-drop, radio buttons allow for excluding and including of variables. Configurations are editable via a graphical dialog. A particular focus was laid on fast SVG rendering by mapping `MooViE` elements to appropriate SVG data types. For instance, curved lines for chords and polylines are mapped to custom Bézier curve approximations, rather than a multitude of line segments.

The `MooViE` tool is developed on Linux/Unix-based systems using CMake (https://cmake.org). It supports platform-as-a-service (PaaS), via a Docker image shipped with the software, thereby being platform-independent. This was tested for Linux and Windows, with scripts automating the Docker setup for these platforms being available. Documentation along with a gallery and explanatory videos is available at https://moovie.readthedocs.io.

## 3 Application examples

The versatility and expressiveness of `MooViE` scenes in combination with interactive data exploration using the `MooViE` engine is shown for three applications from different domains. Configuration files, data and scripts to reproduce the show cases are available in SI Data.

### 3.1 Application 1

We use `MooViE` to study the interrelations between parameters of an epidemiological model (inputs) and its forecasts (outputs) (SI Text Sec. S.1). Specifically, a sensitivity analysis with a state-of-the-art compartmental model for COVID-19 was performed. Of particular interest are the model parameters that have major impact on the outcomes in terms of predicted death counts over a forecast period spanning several weeks. By visually identifying and interactively excluding insensitive parameters, the effects of uncertainty for most important parameters on the predicted time series are stressed. For example, for low-contact rates, the death counts are comparatively insensitive towards the literature range found for the contact-related transmission probability.

On the other hand, the literature range for the probability of dying given a critical course of COVID-19 has a high impact on the death counts, independent of the actual transmission probabilities. Knowing which parameter ranges the outcome is sensitive to is instrumental for forecasting accuracy and, thus, policy-makers at local health authorities. This example shows the capacity of `MooViE` to visualize input-output relations of a nonlinear ODE model, with time-series data as output variables.

### 3.2 Application 2

To demonstrate the effectiveness of `MooViE` in visualizing complex multivariate data from a domain-specific field of systems biology, we exemplify the explorative and bidirectional decision-making workflow in the context of experimental design (SI Text Sec. S.2). Specifically, a comprehensive experimental design study was performed for isotope-labeling experiments to quantify metabolic reaction rates (fluxes), a field called ^13^C metabolic flux analysis. The design goal is to determine those isotope tracer combinations (inputs) that have the best information-to-cost ratio (outputs). To this end, multi-objective studies were performed with and without prior knowledge about the metabolic fluxes. In both, `MooViE` is used to inform experimenters in identifying good trade-off solutions, which are highly informative, but remain cost-efficient. Due to the complexity of the input-output mapping, this is typically a step-wise interactive exploration process. Once the range of suitable experiments is decided, `MooViE` supports the “backpropagation” of these Pareto solutions back to the design compositions, i.e., the tracer combinations that are to be used in the next experiment. For example, among numerous alternatives, glucose species that are isotopically labeled at the first and last position, are preferable tracers, independent of the analytical platform.

### 3.3 Application 3

Comparing the performance of numerical algorithms for solving a class of problems is a ubiquitous task in computer science. Here, we use `MooViE` to provide visual summaries to compare algorithms (SI Text Sec. S.3). Specifically, different optimization algorithms are employed to solve two classic test problems in the field of multi-objective optimization. `MooViE` scenes provide interpretable information about high-dimensional Pareto fronts and facilitate a quick comparative analysis of solution quality. `MooViE` thereby offers a solution to the challenging task of Pareto solution visualization, summarizing the interrelations of design and objective space holistically. Such summaries may be automatically generated within benchmark workflows using `MooViE`’s CLI interface.

## 4 Conclusion

`MooViE` is an C++ package implementing an expressive single-view visualization approach for multivariate datasets with input-output semantic, which commonly occur in data mining pipelines, simulation workflows, or experimental design studies. The tool provides extensive customization features that allow domain scientists to interactively explore and comprehend information from their data and to visualize those in high-quality and appealing diagrams. By its defined interfaces and scalable implementation, `MooViE` has the potential to support domain-scientists in analyzing complex datasets and making scientific discoveries.

## Acknowledgements

The authors thank Sebastian Niedenführ for inspiration of the visualization approach and Wolfgang Wiechert for continuous support.

