## Supplementary Text for "MooViE – Engine for single-view visual analysis of multivariate data"

### Contents

**S.1Compartmental Models in Epidemiology** **2**

**S.2Experimental Design for Isotope Tracer Experiments** **5**

**S.3Visual Assessment of Algorithmic Multi-objective Optimizer Performances** **14**

### S.1 Visualizing the Sensitivity of Compartmental Models in Epidemiology with `MooViE`

##### Motivation

Providing reliable epidemiological forecasts is crucial to mitigate potentially harmful consequences for public health and economy, as has been recently shown in the COVID-19 pandemic with to date almost 7 million confirmed deaths worldwide [WHO, 2023]. On the one hand, epidemiological models provide interpretable insights into the "mechanisms" of previous outbreaks, on the other hand, these models are useful for predicting the evolution of the disease. Attempting to describe the spread of infectious diseases on the population level in an increasingly realistic fashion, compartment models have grown in complexity as more and more details are added [Kühn et al., 2023]. Here challenges arise, however, because these increasingly fine-grained models need to be calibrated with noisy, sparse and aggregated data. When faced with a newly built model, it is therefore important to understand how the model predictions are connected to the underlying parameter space.

##### Case Study: Gaining an understanding how parameters affect the total death count

→ **Key point:** Visual sensitivity analysis for time-resolved epidemic forecasting

To exemplify the visualization concept of `MooViE`, we consider the state-of-the art SECIR model, which distinguishes individuals depending on whether they are susceptible (S), exposed (E), i.e., carry the virus, but are not yet infectious to others, carriers (C), who carry the virus and are infectious to others, but do not show symptoms, infected (I), who carry the virus with symptoms and are infectious to others, and recovered (R), who are no longer infectious to others and not susceptible [Kühn et al., 2023]. Since the SECIR model does not include reinfections of recovered individuals, the dead count approaches a stationary *end-point*. The here investigated SECIR model has 10 parameters in total (see Fig. S.1).

As the number of deaths related to infectious diseases is a statistic that is an often discussed in the public, we here focus on the forecasted death count. Specifically, we aim to understand the quantitative relation between the independent model parameters (**input**) and the number of deaths after a time period of 7, 14, 21, 28 days and at the end point ($10^4$ days), respectively (**output**). For each of the 10 model parameters, we extracted the maximum and minimum values from a literature screen for SARS-CoV-2 [Kühn et al., 2021, Koslow et al., 2022]. We then created the Cartesian product of these values to obtain $2^{10}$ points spanning the parameter space. In addition, we considered the mean of each parameter range. In total, this gives $2^{10} + 1 = 1,025$ parameter constellations (**data items**). For each parameter constellation, we simulated the spreading of the disease starting with 10,000,000 (S), 10,000 (E), 1,010 (C), 1535 (I) and 11 (R) until the stationary end-point using MEmilio [Kühn et al., 2023]. Results were saved in a spreadsheet (`S1_A_table.csv`).

#### Visualization

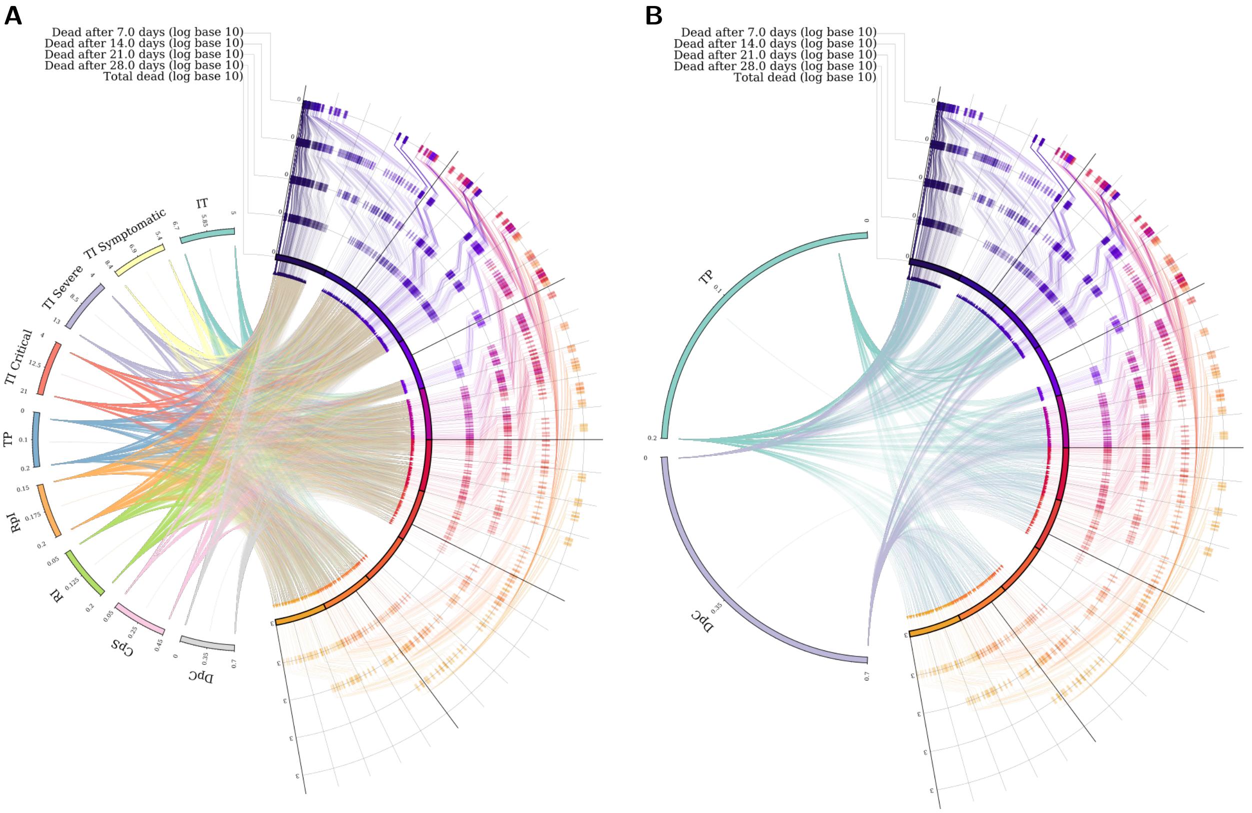


Figure S.1: Two `MooViE` scenes showing influence of parameter combinations on the development of number of deaths over a time span of 1, 2, 3, 4 weeks and in the long run (stationary end-point). **A** shows all 10 varied parameters, and **B** highlights the TP and DpC parameters for the same simulation study as in A. Model parameters: IT – Incubation Time, TI Symptomatic – Time Infected Symptomatic, SI – Serial Interval, TI Severe – Time Infected Severe, TI Critical – Time Infected Critical, TP – Transmission Probability on contact, RpI – Recovered per Infected no symptoms, RI – Risk of Infection from symptomatic, SpIS – Severe per Infected Symptoms, CpS – critical per Severe, DpC – Deaths per Critical. All output arcs are log-scaled and ordered chronologically, with the end-point on the primary arc. High deaths counts are encoded by orange polylines, medium deaths by red and magenta polylines, and low death counts by purple polylines. Histograms for the input samples are turned off, because the distribution of input parameters is not relevant in for this case study.

The `MooViE` scenes in Figure S.1 present an overview of the relationship between predicted deaths over a time course and the reported parameter ranges for the SECIR model. The total number of deaths at the end-point is selected as the primary output arc and the remaining output arcs are arranged in reversed chronological order, in weekly steps. All output arcs are scaled logarithmically and have the same range. The polylines, connecting the death counts between the output arcs, therefore represent the time development of (logarithmic) death counts. In other words, following the polylines in radial direction is equal to following the series of deaths backwards in time. On the input segments, model parameters are arranged, with histograms turned off because their distribution is not relevant for the purpose of our study.

In Fig. S.1 **A**, the result with all parameter constellations is displayed. Focusing on the outputs, i.e., the death count, we see that, strikingly, the range of is quite wide: the lowest simulated total death count is 1 and the highest almost 1000. It is also obvious that the result after 4 weeks is almost identical to the end-point, which is interesting because after 3 weeks we still see a significant increase in deaths for many parameter constellations. By observing the color distribution on the output arcs, we see, that the number of deaths after a certain time strongly correlates with the end-point. The weakest correlation is observed after 1 week, and here we see many parameter constellations, that start off with a rather low number of deaths, but show drastic increases in the subsequent weeks, leading to many crossing polylines. The crossing polylines (as well as the slightly scrambled colors on the output arcs) indicate a change in the development of death counts: some parameter constellations show a high death count initially that remains high, whereas other parameter settings start with a low death count that grows dramatically over time.

To specifically discuss the effect of the parameters TP (transmission probability on contact) and DpC (deaths per critical case), we turned off the other parameters in the `MooViE` scene and arrive at Fig. S.1 **B**. The intuitive connection between high death counts and the DpC parameter is visually confirmed by following the high death count polylines (orange) backwards to the chords into the parameter arcs. None of the high death outcomes (orange) map back to a low DpC parameter value. In contrast, some high DpC values indeed map to lower (purple) death outcomes. By visual comparison, we conclude that the number of deaths is less sensitive to the TP parameter than for the DpC parameter, since both high and low values of the TP parameters lead to high and low death counts. A more accurate prediction of the death count requires an improved estimation of the DpC parameter. Summarizing, in this case study `MooViE` helps to visualize and explore the complex relationships between parameter ranges, as known from literature, and model predictions.

#### S.2 Interpreting the Experimental Design Outcome for Isotope Tracer Experiments with MooViE

##### Motivation

Isotope tracer experiments are used to quantify metabolic rates (fluxes) in living cells [Wiechert and Nöh, 2021]. Since the repertoire of isotope-labeled tracers is vast, with large cost margins, and the tracer experiments are labor-intensive, mathematical models are used to propose informative experimental-analytical settings, a procedure called in silico experimental design [Möllney et al., 1999]. With that, isotope labeling cocktails can be determined by optimizing not just one information criterion, such as the D-criterion [Pukelsheim, 2006], but several, while also taking cost considerations into account [Nöh et al., 2018]. In experimental design, we distinguish between optimal experimental design (O-ED), which is performed at a specific operating point in the flux space using prior knowledge, and robust experimental design (R-ED), which considers all possible fluxes [Beyß et al., 2021].

##### Case Study: Multi-Objective Optimal Experimental Design

→ **Key point:** Visual comparison of complex-structured domain-specific information

In a multi-objective O-ED problem, the objectives defined over a given design space of possible experiments are conflicting. The solution of the optimization problem is therefore not a single solution, but a set of Pareto optimal solutions or Pareto front in the objective space, where a solution cannot be improved in any of the criteria without negatively affecting at least one of the other objectives [Kaisa, 1999]. To interpret the outcome of a multi-objective O-ED problem, means to select a preferred solution on the Pareto front, which is then traced back to its underlying design. The desired functional relationships within the objective and the design spaces, and in particular between both spaces, is obviously highly complex for any but low dimensional problems. For this, only very limited 2/3D visualization techniques are available [Schäpermeier et al., 2021]. We here use `MooViE` to display, navigate and interpret the solution landscape of multi-objective O-ED for the case of tracer experiments.

In our tracer design study, the available tracers are the design parameters. These tracers can be freely mixed (**inputs**). The design is given by a set of labeled species and their fraction in which they are present in the mixture (the sum of the fractions of all contributing tracers adds up to 1). The objectives, on the other hand, are three information criteria (*D-criterion*, *E-criterion*, *A-criterion*), as well as the cost for the experiment (**outputs**). Obviously, for the information criteria high values are preferred, whereas for the cost, low values are desirable.

We performed a multi-objective O-ED for the fungi *Penicilium chrysogenum* growing in a chemostat, comparing two different measurement setups. The `13CFLUX2` simulator [Weitzel et al., 2013] was used to calculate the criterion values for a fine-grained composition grid of the 10 different tracers. Further details on the metabolic model, measurement setups, the cost function, and the computational tool chain used are described in [Nöh et al., 2018]. This information was fed into the multi-objective optimization algorithm SMPSO [Nebro et al., 2009], implemented in the `jMetal` framework [Durillo and Nebro, 2011, Nebro et al., 2015]. In total, 1,000 Pareto optimal results (**data items**) were saved in a spreadsheet (`S2_1_A_table_GCMS.csv` and `S2_1_B_table_LCMSMS.csv`, respectively).

##### Visualization

On the left side of the `MooViE` scenes in Fig. S.2, the 10 glucose tracer species are arranged in segments, on the right side the four objectives are arranged as arcs. As principle arc, we selected the D-criterion, because it is the most commonly used information criterion in the domain-specific field. The chords link the specific designs with their objective values, and are bundled at the principle arc. Histograms indicate the fraction to which the respective tracer species contributes to the mixture. We filtered tracer species that contribute below 1% to a mixture, as these are practically irrelevant.

From the information on the input segments, it immediately becomes clear that some tracer species, e.g., Glc#100001, contribute to many Pareto optimal mixtures, while others, like Glc#010000, are only rarely suggested. When such rare species contribute to a mixture, then only to very low proportion. This

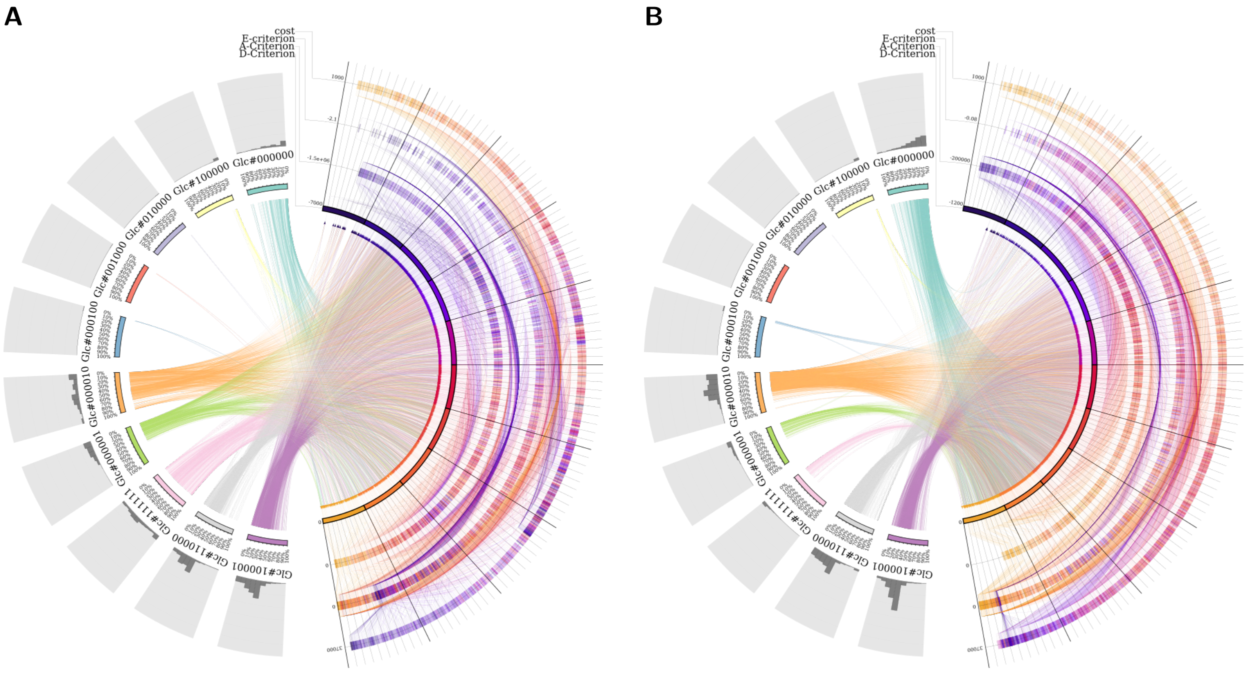


Figure S.2: Pareto optimal mixtures of isotopically labeled glucose (Glc) species. $^{13}$C-labeled positions are indicated by 1, those that are naturally $^{12}$C-labeled by 0. For the D-, A-, and E-criteria higher values are better, for the cost lower values are better, being reflected in the orientation of the arcs. Histograms are showing the fraction with which the tracer contributes to the mixture. **A** – tandem mass spectrometry coupled to liquid chromatography (LC-MSMS); **B** – mass spectrometry coupled to gas chromatography (GC-MS).

information is provided by the histograms and immediately allows making informed decisions on which tracers are relevant. On the right side, the arcs are ordered from primary to secondary importance. In this particular case, information criteria and costs hold the inner and outer arcs, respectively. Moreover, the ranges of the arcs are aligned from preferable (upper end) to undesirable (lower end).

The chords are bundling the single Pareto mixture designs to meet at their corresponding D-criterion value. The specific D-criterion value determines the color of the boxes and the polyline linking the objective values. This allows to spot (anti)correlations between the objectives. For example, D-criterion values and costs are strongly negatively correlated, as the colors on the D-criterion arc appear flipped on the cost arc. This means highly informative designs in terms of the D-criterion are expensive, while the cheapest designs are not informative in terms of the D-criterion. With `MooViE`, we can work with the scenes interactively, so that the ordering of the arcs can be changed on-the-fly. This is useful, e.g., when some objectives turn out to correlate and reordering the arcs rectifies the polylines.

For Fig. S.2, we used `MooViE` to compare the outcome of two in silico designs, one for tandem mass spectrometry coupled to liquid chromatography (LC-MSMS, Fig. S.2 A) and one for mass spectrometry coupled to gas chromatography (GC-MS, Fig. S.2 B). When comparing the outcome of the two different measurement setups, we can see that the same tracer species are proposed, but to different fractions. For instance, Glc#100001 is an important tracer species in both cases, but with GC-MS it is more often chosen. Glc#000010 and Glc#111111, on the other hand, are only selected with fractions below 50% for GC-MS, while for LC-MSMS fractions of up to 100% appear.

Comparing D-, A-, and E-criteria values in absolute numbers is generally not reasonable, but comparing the generated `MooViE` scenes gives insights into their correlations. For instance, whereas the D- and cost values are almost perfectly anti-correlated for GC-MS as explained above, for LC-MSMS the outcome is more diverse. Indeed, mixtures are proposed that achieve a good trade-off between the two criteria. This is visible from the numerous orange-yellow boxes (indicating relatively high D-criterion values) in the mid-range of the cost arc. How to interactively further distill information about these specific mixtures is described in the next case study.

Despite the domain-specificity of this application, the kind of visualization provided by `MooViE` is transferable to experimental design in other fields, and more generally any multi-objective optimization problem. With that `MooViE`, may be a trailblazer for a new way of visualization in this field (see also Sec. S.3).

**Case Study: Robust Multi-Objective Experimental Design**

→ **Key point:** Interactive visual exploration and thinning workflow

In a related design content, rather than looking for the Pareto mixtures at a specific operating point of a system (flux constellation), we want to explore solutions for any possible operating point (flux space). For example, in a first-of tracer experiment, cheap mixtures with few components that are nevertheless considered sufficiently informative with respect to a sub-set of key parameters may be preferred. Such R-ED thus adds another layer of complexity for the interpretation of ED results, which makes the interactive and exploratory examination of the results necessary.

The input are the same as in the first case study (tracer species), while for R-ED additional objectives come into play. One new metric, called coverage ($\Phi_{cover}$), is concerned with the proportion of the flux constellations for which a design is informative at all. Other R-ED objectives are concerned with the average D-criterion ($\Phi_{\bar{D}}$). We previously performed R-ED for *Streptomyces clavuligerus* [Beyß et al., 2021] for two tracers, glycerol (Glyc) and arginine (ARG). The amount of Glyc and ARG was fixed for the computations. Again, the `13CFLUX2` simulator was used to calculate the values of 4 criteria (**outputs**) for 7 tracer species (with fractions gridded in steps of 10%) for 1,000 samples of the parameter space (**inputs**). For more details about the computational workflow and an in-depth explanation of the metrics, we refer to Beyß et al. [2021]. As a result of the calculations, in total 18,776 data items were saved in a spreadsheet (`S2_2_table_RED.csv`).

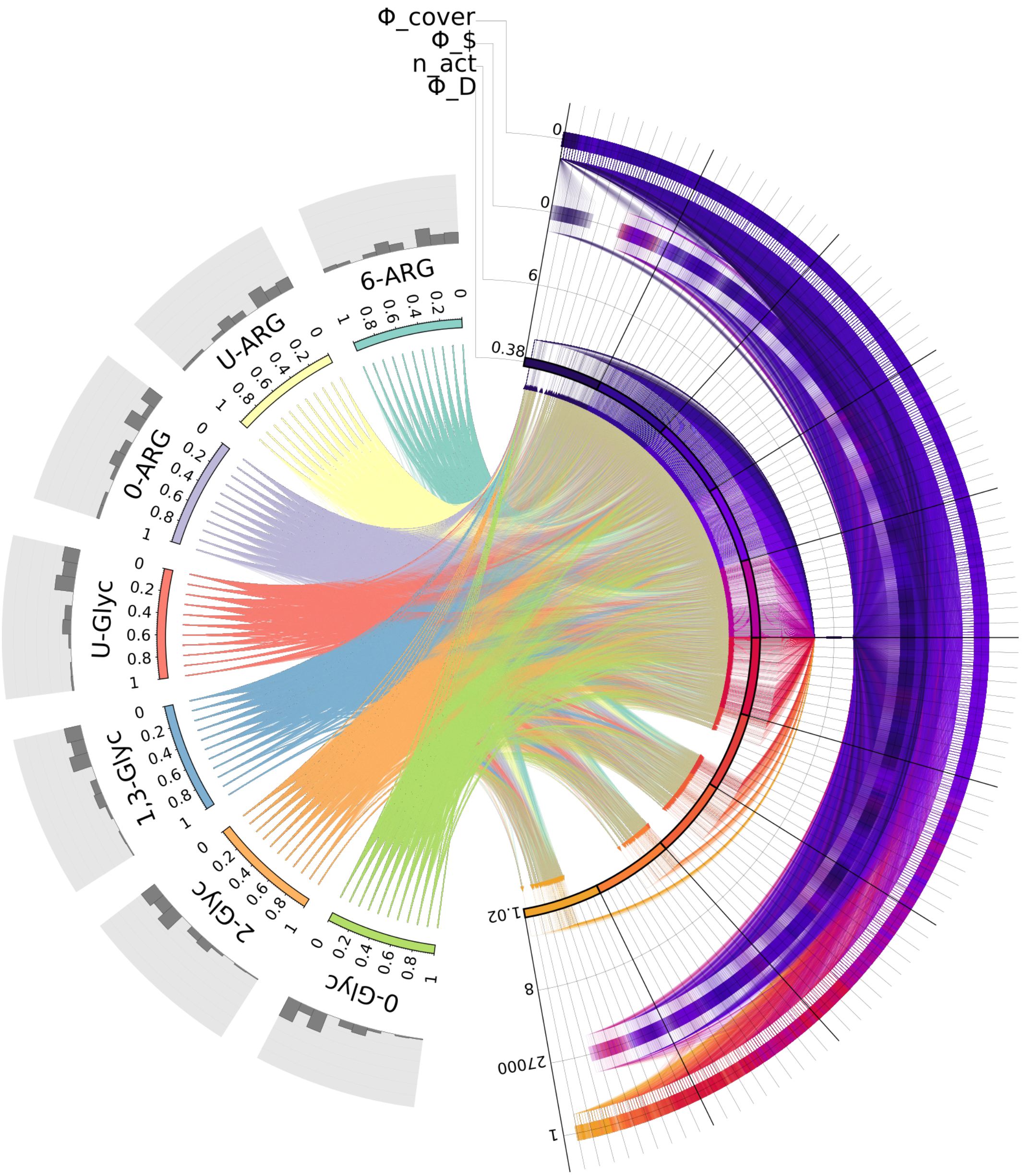


Figure S.3: `MooViE` scene showing mixtures of isotopically labeled glycerol (Glyc) and arginine (ARG) species (input) evaluated for several criteria (output). Numbers before the substrate name indicate the position of $^{13}$C. For all criteria, higher values are better. The image comprises almost 20,000 data-points. The nuisance parameter n_act is identical for all data-points, reducing the traceability of the polylines, thereby hampering interpretation.

#### Visualization

Figure S.3 shows the rendered `MooViE` scene. The vector graphic is over 100MB large and consists of almost 300,000 path elements. `MooViE` reduces the processing burden of displaying the images by restricting to the SVG Tiny subset [W3C, 2008]. With this, even such comprehensive scenes can still fluently created using `MooViE`. However, the interactive handling of such comprehensive scenes starts to become somewhat impractical.

Again, on the left side the tracer species are arranged, the right side shows the objective values. With its 18,776 designs, the `MooViE` scene is very crowded. It becomes clear by the colors that the coverage metric

($\Phi\_cover$) on the inner arc, strongly correlates with the average D-criterion ($\Phi_{\bar{D}}$) on the outermost arc. However, further visual interpretation is hampered by a nuisance variable displayed on the second inner arc, where the polylines of all designs meet at one value.

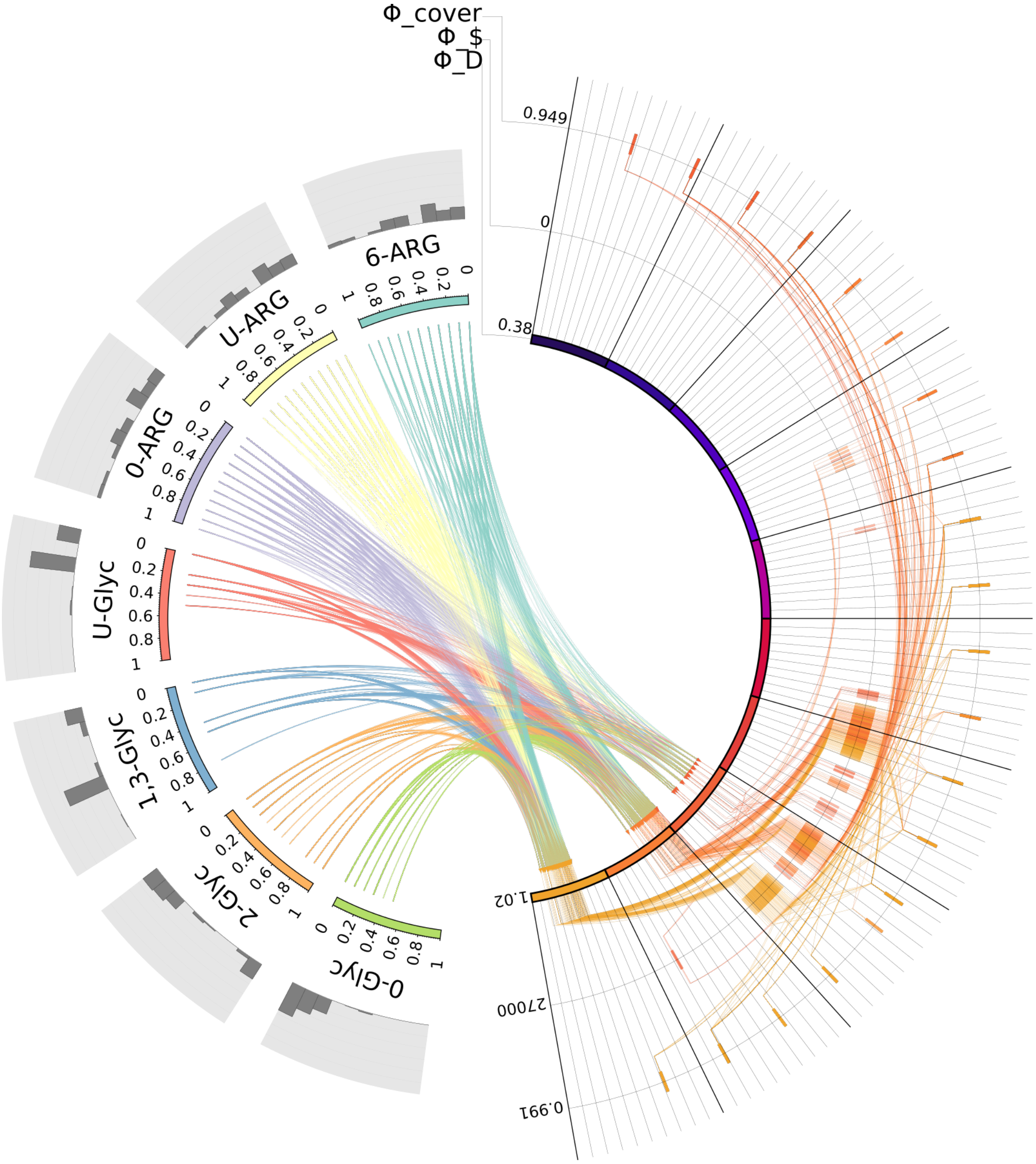


Figure S.4: `MooViE` scene after eliminating $n_{act}$ and limiting $\Phi\_cover$ to reasonably high values.

For identifying a good tracer mixture, this arc is removed from the `MooViE` scene. Naturally, we are only interested in design, with a high probability for being informative. so, as in [Beyß et al., 2021], we constrained the coverage by $\Phi\_cover \geq 0.95$. This allows exploring the relationships hidden in the data in more detail. The result in Fig. S.4 comprises the remaining 287 mixtures. Here it becomes visible that the coverage actually takes discrete values, as it refers to a fraction of parameter constellations. The average D-criterion ($\Phi_{\bar{D}}$) and the cost metric ($\Phi_{\$}$), are not well scaled. We therefore set cost limits to $0.82 \leq \Phi_{\bar{D}} \leq 1.02$ and $9,000 \leq \Phi_{\$} \leq 22,000$. By this, the two most expensive mixtures are eliminated. Both mixtures have relatively low D-criterion and coverage values, as visible from the polylines.

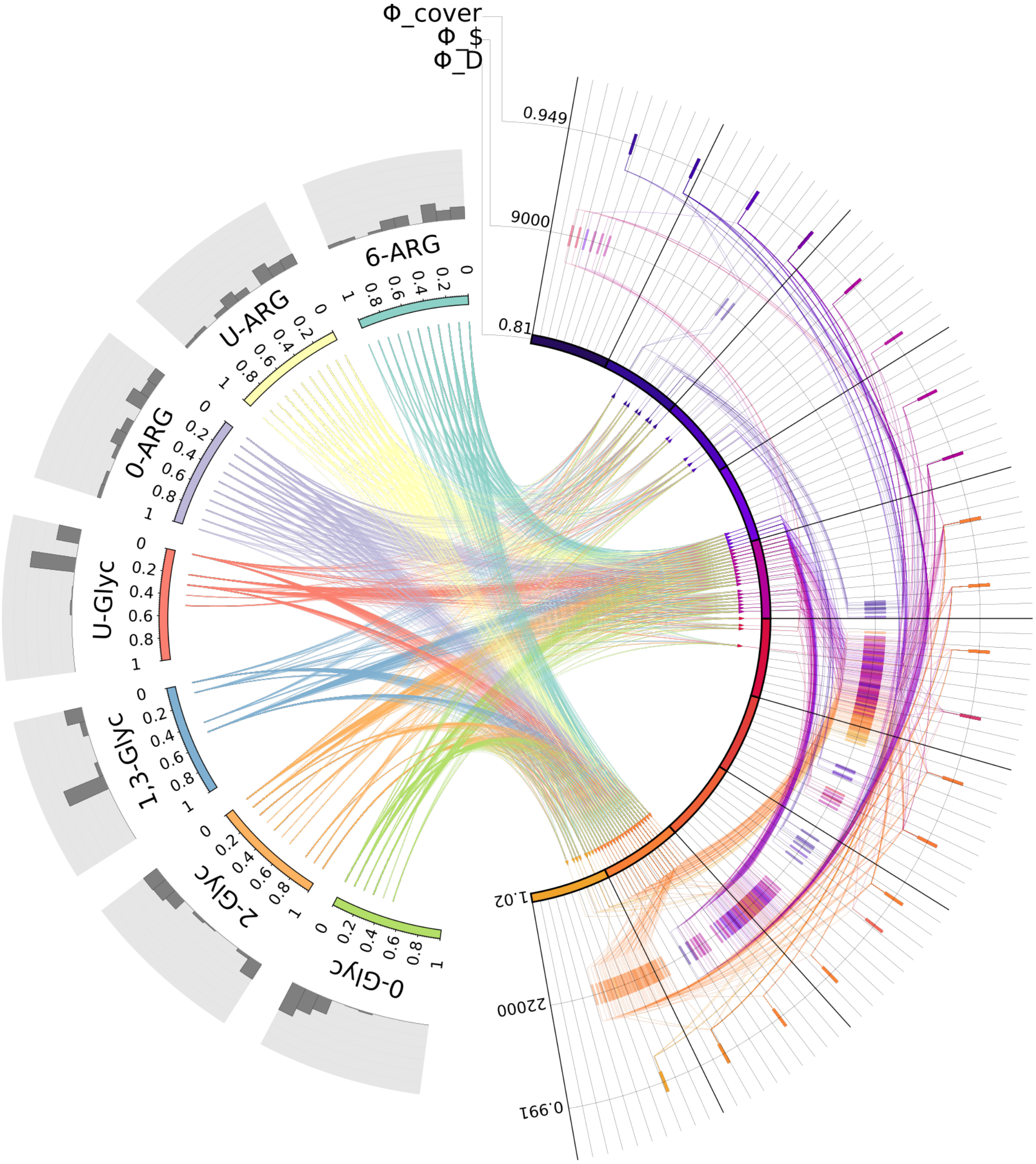


Figure S.5: `MooViE` scene after rescaling $\Phi_{\bar{D}}$ and $\Phi_{\$}$ and eliminating two overly costly mixtures.

After elimination, 285 mixtures remain in Fig. S.5. On the cost arc, we spot a rather interesting region, in the mid-price segment, with highly informative designs that are still affordable. We select these designs, by further constraining the cost margin to $15,500 \leq \Phi_{\$} \leq 17,000$, leaving 132 mixtures in Fig. S.6.

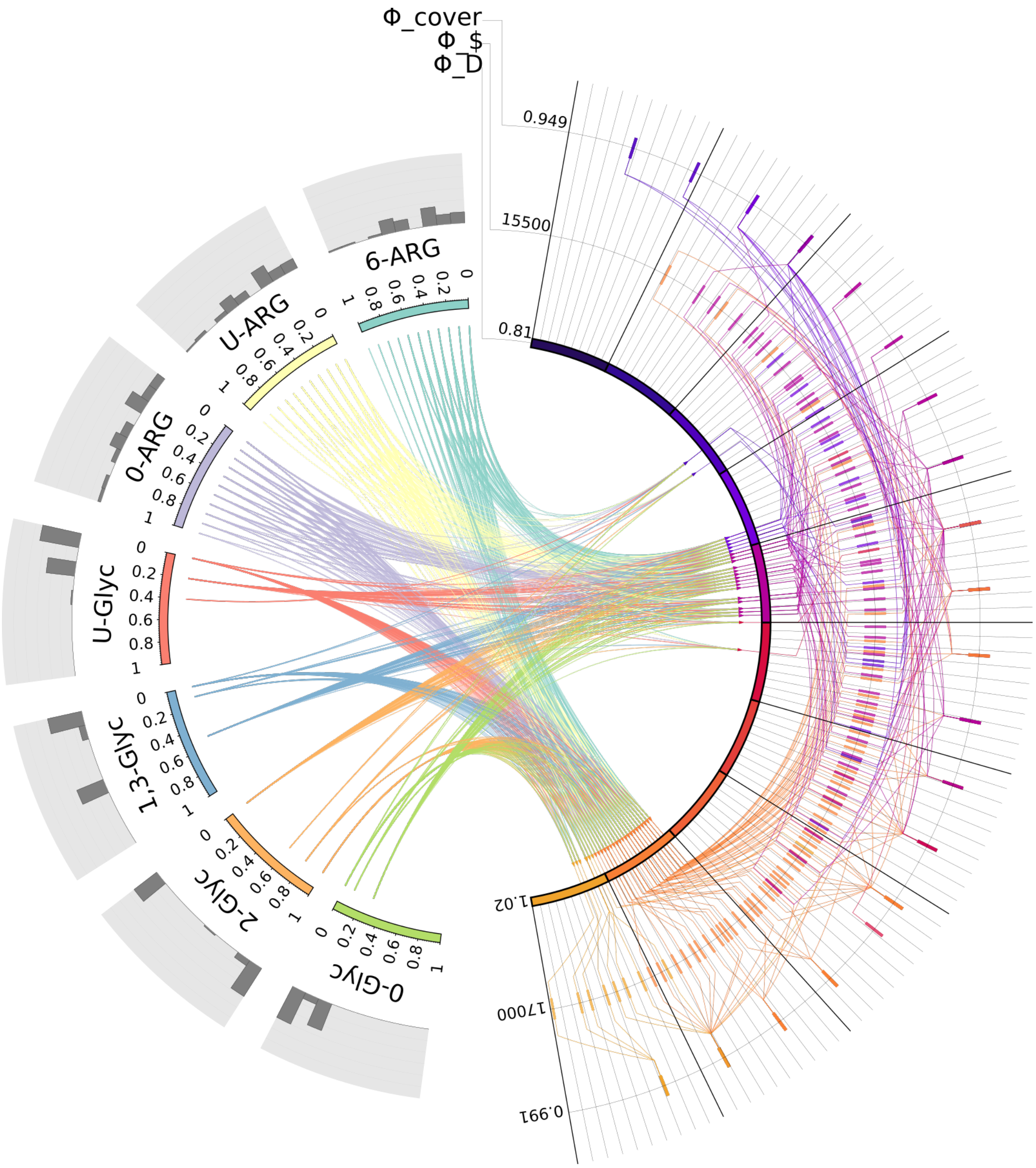


Figure S.6: `MooViE` scene after zooming in on affordable, yet informative designs.

Like before, we see that for the Glyc mixtures many combinations are excluded, while for all ARG species, there are still many connecting arcs for all labeling fractions. From this we concluded that, compared to Glyc, ARG mixtures are only of minor importance. Therefore, we excluded the latter from the scene for clarity. This is one of the scientific findings reported in the original publication. For readability, we finally re-scaled the D-criterion to be $\geq 0.86$ (cf. Fig. S.7).

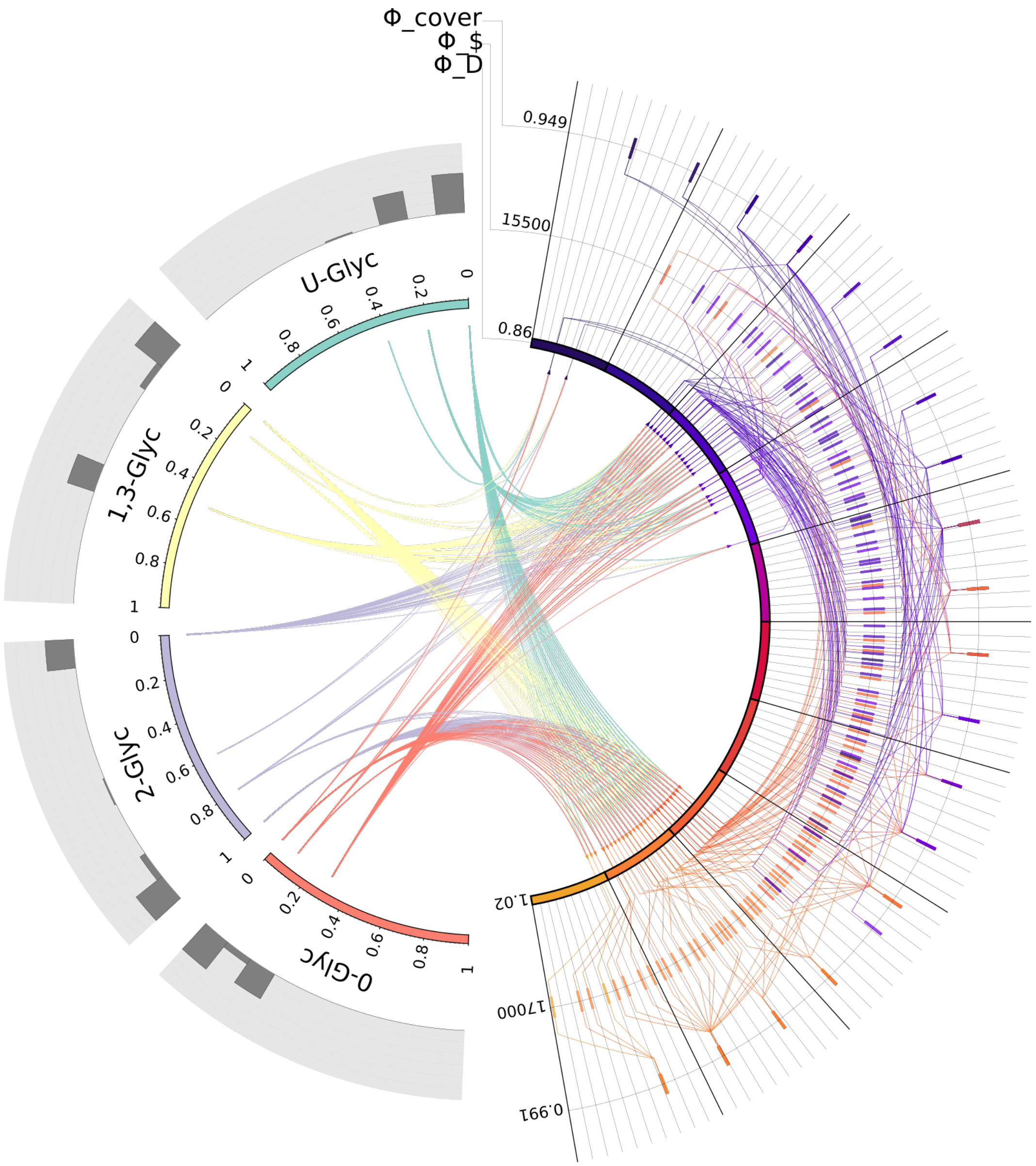


Figure S.7: `MooViE` scene after excluding the non-informative ARG mixtures, due to their minor practical importance, and rescaling the D-criterion.

With regard to the average D-criterion, the mixtures fall in two categories: (1) D-criterion values of approximately 1, and (2) D-criterion values of approximately 0.9. Both cases exhibit very specific Glyc mixtures. For the first case, 2-Glyc is dominant, with fractions of 100% or 80%. The remaining percentages are filled by 0-Glyc and 1,3-Glyc. For the second case, 1,3-Glyc is mixed with U-Glyc and 0-Glyc, with 1,3-Glyc mostly being 50%. When turning to the costs, it is important to keep in mind that we have already limited the cost margins, so the most expensive designs are not more than 13% more costly than the cheapest ones. While the majority of the mixtures are on the upper end of the price range, there are still plenty of designs at the lower range, for example, in case no 2-Glyc is used, but instead 50% 1,3-Glyc, 30% 0-Glyc and 20% U-Glyc. Concerning the costs, these mixtures are on average cheaper. From these considerations, we selected two viable candidates for highly informative, yet affordable mixtures. One is the pure 100% 2-Glyc, whereas the other suggested mixtures, of 50% 1,3-Glyc and 30% 0-Glyc and 20% U-Glyc, was identified by putting higher importance to the cost.

In summary, `MooViE` allowed us to screen and zoom in on interesting regions of the output space. Thereby, we uncovered the insensitivity of the objectives with respect to the ARG labeling. in contrast, high proportions of the positionally labeled 1,3-Glyc or 2-Glyc tracers are mandated for performing informative experiments.

### S.3 Comparative Visual Assessment of the Solution Performance of Multi-Objective Optimization Algorithms with MooViE

##### Motivation

In numerical mathematics, many algorithms exist that tackle the same problem formulation, but employ conceptually different solution strategies. Because exhaustive comparisons are often infeasible for real-world problems, lower-dimensional synthetic test problem with known analytic solutions are employed. We here sketch this procedure using an example from multi-objective optimization (MOO), which for instance occurs in experimental design (see Sec. S.2).

MOO problems have the general form

$$\min_{x \in X} (f_1(x), \dots, f_n(x)) \tag{1}$$

where $X \subseteq \mathbb{R}^m$ is the *design space* and $Y = \operatorname{img}(f_1, \dots f_n) \subseteq \mathbb{R}^n$ the *objective space*. Typically, these problems have many trade-off solutions. These trade-off solutions consist of all designs for which the vector of objectives cannot be improved in any single objective without degrading in another. The design vectors are called *Pareto-optimal set* $X^* \subset X$ and the respective objective values $Y^* = \operatorname{img}(f_1|_{X^*}, \dots f_n|_{X^*})$ *Pareto front*. The shapes of the Pareto fronts, whose characterization is the analysis goal, can take various forms: they may be regular convex, describe a sub-dimensional manifold, or may even be discontinuous (see Fig. S.8).

Real-world MOO problems in Eq. (1) are analytically intractable. Thus, Pareto-optimal sets and Pareto fronts have to be approximated numerically. For this, a plethora of multi-objective algorithms have been developed, differing in their strategies to explore the design space [Pereira et al., 2022]. The solution performance, in particular with regard to the coverage of the design space and Pareto front, provided by these algorithms is therefore compared by means of synthetic test problems with at least partially known solutions [Deb, 2001]. We here show that MooViE scenes are an effective way to survey the characteristics of Pareto solutions with their intricate relations between Pareto sets and fronts (Case study 1), as well as to qualitatively compare the outcome of numerical optimizers (Case study 2).

#### Case study 1

→ **Key point:** Visual summary of multidimensional Pareto set-front relations

For illustration purposes, we took two test problems, namely DTLZ5 and DTLZ7, from Deb [2001], with solutions that exhibit distinct characteristics. For both problems, the design spaces are the four-dimensional hypercube ($0 \leq x_i \leq 1, i = 1, \dots, 4$), and the objective spaces are three-dimensional ($f_i, i = 1, \dots, 3$). For

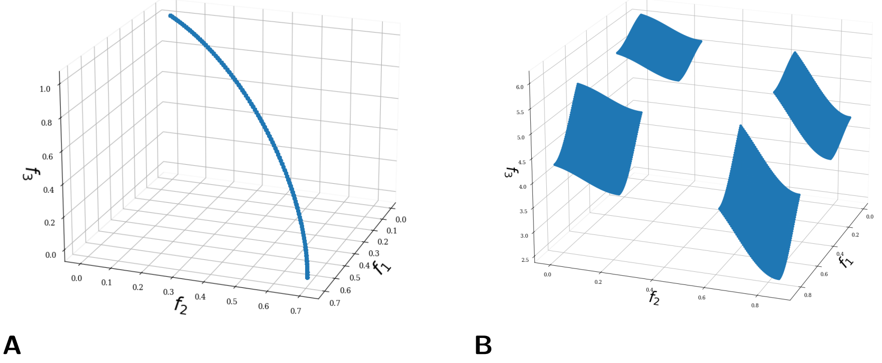


Figure S.8: Pareto frontiers of test problems DTLZ5 (**A**) and DTLZ7 (**B**). The objective values are aligned on a continuous two-dimensional manifold and four discontinuous square-shaped patches, respectively.

the test problem DTLZ5, the Pareto set is known to fulfill $x_3 = \frac{1}{2}, x_4 = \frac{1}{2}$, and the Pareto front is constrained by $\sum_{i=1}^{4} f_i^2(x) = 1$ (see Fig. S.8 **A**). DTLZ7, on the other hand, has a discontinuous Pareto front that consists of four disjoint "patches" (see Fig. S.8 **B**), and its Pareto set satisfies $x_3 = 0, x_4 = 0$. Pareto solutions of both test problems, DTLZ5 and DTLZ7, were calculated, using the well-known NSGA-II algorithm implemented in pymoo Deb et al. [2002], Blank and Deb [2020]. In each case, a solution set of 100 designs (**inputs**) and associated objective values (**outputs**) was obtained (**data items**). The results are available as spreadsheets S3_1_table_DTLZ5.csv and S3_1_table_DTLZ7.csv.

##### Visualization

In the MooViE scene shown in Fig. S.9, design variables are arranged on the input side on the left of the scene, whereas the objectives are presented on the output side on the right. The output arcs are arranged according to the indexing of the objective functions, from 1 (primary arc) to 3 (outer arc). Likewise, input segments, representing the design values, are ordered according to their index (1 on the top to 4 at the bottom). Histograms associated with the input arcs display the distribution of calculated designs among the data items.

The solution for test problem DTLZ5 is shown in Fig. S.9 **A**. Starting with the designs, we observe that NSGA-II found the correct solution for the design variables $x_3$ and $x_4$ ($\frac{1}{2}$) as indicated by the histograms (red arrows). The $x_1$ design variables, on the other hand, are spread over their range ($[0, 1]$), whereas those for $x_2$ are predominately located close to 1, but also to a minor degree between 0.1 and 0.5. Starting from the input segments, the chords connect to their associated $f_1$ value located on the primary output arc, taking values between 0 and 0.7. The chords show that $f_1$ is inversely correlated to $x_1$, i.e., low (high) values of $x_1$ map to high (low) values of $f_1$. For the other design variables, no such statement can be made, implying that their relation to $f_1$ is governed by a specific composition of values. For the output arcs, we see that the value range for $f_1$ is almost uniformly covered. Parallel polylines and their colors indicate that $f_1$ and $f_2$ align, while a crossing of polylines and their color flip connecting $f_2$ and $f_3$ values in a single point (red arrow) conveys that on the Pareto front $f_3$ is strictly decreasing in $f_2$. These observations on the continuity and correlations of the Pareto front is validated by comparing the MooViE scene to the true Pareto front in Fig. S.8 **A**. The fact that also the value range of the $f_3$ arc is more or less evenly covered with no color flips indicates the continuity of the Pareto front for DTLZ5.

Compared to this, the polylines of the MooViE scene for the second test problem, DTLZ7, are distinctly different. Here, in Fig. S.9 **B** large gaps in the coverage of the output arcs are explained by the discontinuity of the Pareto front (see Fig. S.8 **B**, two-sided red arrow). Again, the histograms for the input variables $x_3$ and $x_4$ indicate that the true solution (0) was found (red arrows). On the input segments of $x_1$ and $x_2$, the histograms identify the values that are more frequently contained in the Pareto set determined by NSGA-II. For both design variables, these values are located at the higher and the lower ends of their value ranges. Compared to the DTLZ5 scene, the characteristics of the chords starkly differ, reflecting the differences in the Pareto frontiers in Fig. S.8.

**A**

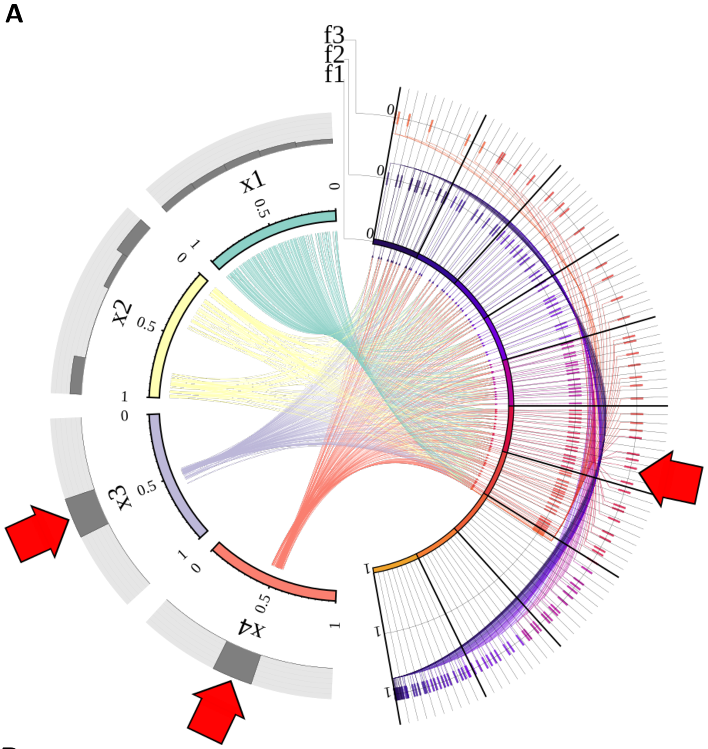


**B**

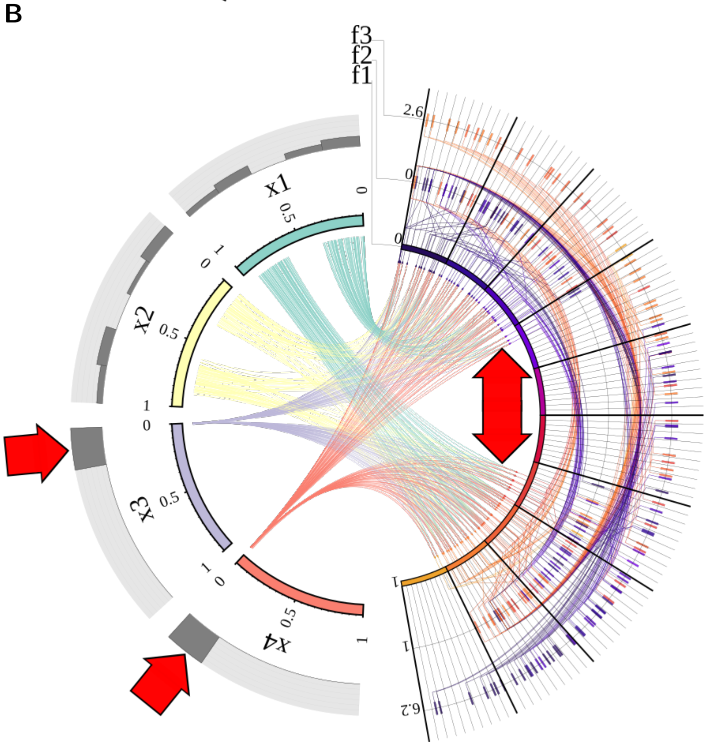


Figure S.9: `MooViE` scenes capturing the numerical Pareto solutions of test problems `DTLZ5` (**A**) and `DTLZ7` (**B**). Input segments represent design variables, output arcs represent objective values. Arrows point to specific characteristics of the Pareto solutions.

#### Case study 2

→ **Key point:** Visual comparison of numerical Pareto set-front relations

In the last case study, for solving the two test problems, a single algorithm (NSGA-II) was used. This algorithm belongs to the class of multi-objective evolutionary algorithms (MOEA), which approximate the Pareto front by evolving a set of particles using selection operators. Different to the selection operator employed in NSGA-II, other algorithms employ so-called indicator- or decomposition-based selection strategies [Emmerich and Deutz, 2018]. We now use MooViE scenes to visually compare Pareto solutions produced using algorithms that employ different selection strategies.

Again, Pareto solutions of the two test problems, DTLZ5 and DTLZ7, were computed. Besides NSGA-II, we used two additional algorithms, namely SMS-EMOA and MOEA/D, which employ indicator- and decomposition-based strategies, respectively. Both algorithms are implemented in pymoo. For each test problem and each MOO algorithm, a solution set of 100 designs (**inputs**) and associated objective values (**outputs**) was obtained (**data items**). The results were collected in spreadsheets that follow the pattern `S3_2_table_<problem name>_<algorithm name>.csv`.

##### Visualization

For each of the six problem-algorithm combinations, one MooViE scene in Fig. S.10 visualizes the computed numerical solution. For direct comparability, the order and ranges of all input and output arcs are kept the same across algorithms, as described before.

To compare the solutions, we start with looking at the first column in Fig. S.10, showing the solutions for test problem DTLZ5. All three algorithms, NSGA-II, SMS-EMOA and MOEA/D, capture the basic characteristic properties of the true Pareto solution, as discussed in Case study 1 (red arrows in Fig. S.9). However, the visual comparison of the three MooViE scenes immediately reveals that the MOEA/D scene considerably differs from the two others: MOEA/D resolves the Pareto front at distinct points only, in contrast to the other algorithms that provide a much finer resolution. This is also indicated by the polylines, which only connect distinct values on the output arcs for MOEA/D, although covering the ranges of the true solution. SMS-EMOA provides solutions closer to those of NSGA-II. Nonetheless, as the histograms show, the determined Pareto sets still differ in the distribution of $x_1$ values and the coverage of $x_2$, for which SMS-EMOA covers almost the full range between 0 and 1.

Turning to the solutions for the second test problem DTLZ7 in Fig. S.10, we find that the basic characteristic properties of the Pareto solutions are largely matched by all algorithms. The MooViE scene for MOEA/D reveals, however, that the "patches" of the Pareto front (see Fig. S.8 **B**) are determined in an unbalanced way. Here, only very few points of the "patches" associated with low values of $x_1$ or $x_2$ are contained in the solution set. With regard to the input arcs and chords, we see that MOEA/D especially covers the low ranges of $x_1$ and $x_2$ worse than NSGA-II and SMS-EMOA, leading to poor coverage of the lower ranges of $f_1$.

In summary, MooViE allows quick assessment and comparison of the coverage of Pareto set and Pareto front for both test problems by drawing on their known properties. We believe that MooViE's ability to provide visual summaries is a considerable aid for MOO solution communication in higher dimensional problems, and thereby useful to visual algorithm comparison. We want to emphasize that the comparison here is not to be understood in terms of a representative benchmark of the three MOO algorithms. This example is not the only way MOO algorithms can be compared. A more fine-grained approach may involve a grid of different hyperparameter settings plotted against multiple quality metrics on the results. Here, a visual summary may be equally useful to overview algorithm performance or to provide insights on effectiveness and robustness of tuning. Clearly, this approach is also transferable to any other numerical algorithm.

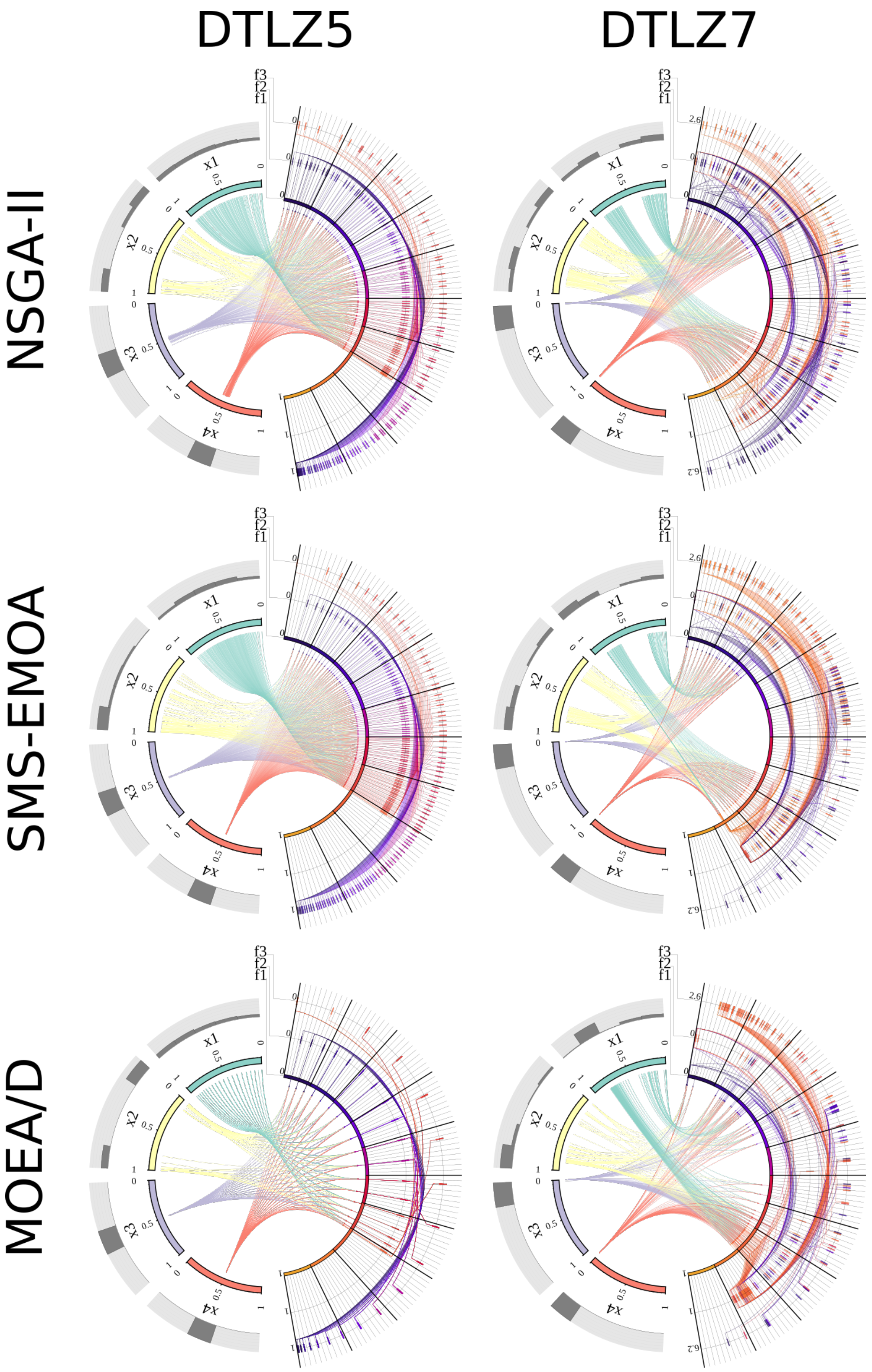


Figure S.10: `MooViE` scenes displaying the solutions of the numerical test problems `DTLZ5` and `DTLZ7` (columns), computed by the algorithms `NSGA-II` (see also Fig. S.9), `SMS-EMOA` and `MOEA/D` (rows). Input segments and output arcs represent design variables and objectives, respectively.

Kalyanmoy Deb. *Multi-Objective Optimization Using Evolutionary Algorithms*. Wiley-Interscience Series in Systems and Optimization. John Wiley & Sons, 1st edition, 2001.

Kalyanmoy. Deb, Amrit Pratap, Sameer Agarwal, and T. Meyarivan. A fast and elitist multiobjective genetic algorithm: NSGA-II. *IEEE Transactions on Evolutionary Computation*, 6(2):182–197, 2002. doi: 10.1109/4235.996017.

Juan J. Durillo and Antonio J. Nebro. jmetal: A Java framework for multi-objective optimization. *Advances in Engineering Software*, 42(10):760–771, 2011. ISSN 0965-9978. doi: https://doi.org/10.1016/j.advengsoft.2011.05.014.

Michael T. M. Emmerich and André H. Deutz. A tutorial on multiobjective optimization: Fundamentals and evolutionary methods. *Natural Computing*, 17(3):585–609, 2018. doi: 10.1007/s11047-018-9685-y.

Miettinen Kaisa. *Nonlinear Multiobjective Optimization*, volume 12 of *International Series in Operations Research & Management Science*. Kluwer Academic Publishers, Boston, USA, 1999.

Wadim Koslow, Martin J. Kühn, Sebastian Binder, Margrit Klitz, Daniel Abele, Achim Basermann, and Michael Meyer-Hermann. Appropriate relaxation of non-pharmaceutical interventions minimizes the risk of a resurgence in SARS-CoV-2 infections in spite of the Delta variant. *PLOS Computational Biology*, 18 (5):e1010054, 2022. doi: 10.1371/journal.pcbi.1010054.

Martin J. Kühn, Daniel Abele, Tanmay Mitra, Wadim Koslow, Majid Abedi, Kathrin Rack, Martin Siggel, Sahamoddin Khailaie, Margrit Klitz, Sebastian Binder, Luca Spataro, Jonas Gilg, Jan Kleinert, Matthias Häberle, Lena Plötzke, Christoph D. Spinner, Melanie Stecher, Xiao Xiang Zhu, Achim Basermann, and Michael Meyer-Hermann. Assessment of effective mitigation and prediction of the spread of SARS-CoV-2 in Germany using demographic information and spatial resolution. *Mathematical Biosciences*, 339:108648, 2021. doi: 10.1016/j.mbs.2021.108648.

Martin J. Kühn, Daniel Abele, David Kerkmann, Sascha A. Korf, Henrik Zunker, Anna C. Wendler, Julia Bicker, Dang Khoa Nguyen, René Schmieding, Lena Plötzke, Patrick Lenz, Maximilian F. Betz, Carlotta Gerstein, Agatha Schmidt, Paul Johannssen, Margrit Klitz, Sebastian Binder, Martin Siggel, Wadim Koslow, Jan Kleinert, Kathrin Rack, Annette Lutz, and Michael Meyer-Hermann. Memilio v1.0.0 - a high performance modular epidemics simulation software, 2023. URL `https://elib.dlr.de/201660/`.

Michael Möllney, Wolfgang Wiechert, Dirk Kownatzki, and Albert A. de Graaf. Bidirectional reaction steps in metabolic networks. IV. Optimal design of isotopomer labeling experiments. *Biotechnology and Bioengineering*, 66(2):86–103, 1999. doi: 10.1002/(SICI)1097-0290(1999)66:2⟨86::AID-BIT2⟩3.0.CO;2-A.

Antonio J. Nebro, Juan José. Durillo, Jose Garcia-Nieto, Carlos A. Coello Coello, Francisco Luna, and Enrique Alba. SMPSO: A new PSO-based metaheuristic for multi-objective optimization. In *2009 IEEE Symposium on Computational Intelligence in Multi-Criteria Decision-Making*, number 2, pages 66–73, 2009. doi: 10.1109/MCDM.2009.4938830.

Antonio J. Nebro, Juan J. Durillo, and Matthieu Vergne. Redesigning the jMetal multi-objective optimization framework. In *Proceedings of the Companion Publication of the 2015 Annual Conference on Genetic and Evolutionary Computation*, pages 1093–1100, New York, NY, USA, 2015. ACM. doi: 10.1145/2739482.2768462.

Katharina Nöh, Sebastian Niedenführ, Martin Beyß, and Wolfgang Wiechert. A Pareto approach to resolve the conflict between information gain and experimental costs: Multiple-criteria design of carbon labeling experiments. *PLoS Computational Biology*, 14(10):e1006533, oct 2018. doi: 10.1371/journal.pcbi.1006533.

João Luiz Junho Pereira, Guilherme Antônio Oliver, Matheus Brendon Francisco, Sebastião Simões Cunha, and Guilherme Ferreira Gomes. A review of multi-objective optimization: Methods and algorithms in mechanical engineering problems. *Archives of Computational Methods in Engineering*, 29(4):2285–2308, 2022. doi: 10.1007/s11831-021-09663-x.

Friedrich Pukelsheim. *Optimal Design of Experiments*. SIAM, 2006. ISBN 978-0-89871-604-7.

Lennart Schäpermeier, Christian Grimme, and Pascal Kerschke. To boldly show what no one has seen before: A dashboard for visualizing multi-objective landscapes. In *Evolutionary Multi-Criterion Optimization*, pages 632–644. Springer International Publishing, 2021. doi: 10.1007/978-3-030-72062-9_50.

W3C. Scalable vector graphics (svg) tiny 1.2 specification, 2008. URL `http://www.w3.org/TR/2008/REC-SVGTiny12-20081222/`. Accessed: 2024-02-07.

Michael Weitzel, Katharina Nöh, Tolga Dalman, Sebastian Niedenführ, Birgit Stute, and Wolfgang Wiechert. 13CFLUX2 - High-performance software suite for $^{13}$C-metabolic flux analysis. *Bioinformatics*, 29(1):143–145, 2013. doi: 10.1093/bioinformatics/bts646.

WHO. Coronavirus disease (COVID-19), 2023. Accessed: 2023-10-28.

Wolfgang Wiechert and Katharina Nöh. Quantitative metabolic flux analysis based on isotope labeling. In Jens; Nielsen, Gregory; Stephanopoulos, and Sang Yup; Lee, editors, *Metabolic Engineering: Concepts and Applications*, chapter 3, pages 73–136. Wiley, 2021. doi: 10.1002/9783527823468.ch3.
