## Supplementary material for "MooViE – Engine for single-view visual analysis of multivariate data": README.md

# S1: Compartmental Models in Epidemiology

To simulate data, some dependencies are required.
Fetch the repository https://codebase.helmholtz.cloud/loki/memiliflow
with the commit 8ee5d03e9dc47c5d0ed89af10283c36afe0ffbc6.

Then go into the repository and call:
```
./install_memilio.sh -c "$(<.memilio_commit_hash)"
```

This script installs the python API for memilio version 1.0, see https://doi.org/10.5281/zenodo.10412635. Once this is installed, the data can be simulated by

```
python3 sim_data_secir.py
```

Once the data has been generated (or the existing data is used), the MooViE figures are generable by executing run.sh
