## Supplementary material for "MooViE – Engine for single-view visual analysis of multivariate data": sim_data_secir.py

```
import numpy as np
import pandas as pd
from memilio.simulation import Damping, TimeSeries, secir
from memilio.simulation import ContactMatrix, Damping, UncertainContactMatrix
import itertools
from memilio.simulation import AgeGroup

if __name__ == "__main__":
    num_age_groups = 1
    model = secir.Model(num_age_groups)

    A0 = AgeGroup(0)
    age_groups = [A0]
    # A1 = age_groups[1]
    # A2 = age_groups[2]
    # A3 = age_groups[3]
    # A4 = age_groups[4]
    # A5 = age_groups[5]

    ################################################################################
    # Initial State
    ################################################################################
    model.populations[A0, secir.InfectionState.Susceptible] = 10_000_000
    model.populations[A0, secir.InfectionState.Exposed] = 10_000
    model.populations[A0, secir.InfectionState.InfectedNoSymptoms] = 1_000
    model.populations[A0, secir.InfectionState.InfectedNoSymptomsConfirmed] = 10
    model.populations[A0, secir.InfectionState.InfectedSymptoms] = 1_000
    model.populations[A0, secir.InfectionState.InfectedSymptomsConfirmed] = 500
    model.populations[A0, secir.InfectionState.InfectedSevere] = 30
    model.populations[A0, secir.InfectionState.InfectedCritical] = 5
    model.populations[A0, secir.InfectionState.Recovered] = 10
    model.populations[A0, secir.InfectionState.Dead] = 1

    #serial_interval = [
        #0.5 * 2.67 + 0.5 * 5.2,
        #(0.5 * 2.67 + 0.5 * 5.2 + 0.5 * 4.00 + 0.5 * 5.2) / 2,
        #0.5 * 4.00 + 0.5 * 5.2,
    #]
    # time_infected_severe = [4., (4. + 13.) / 2., 13.]
    # time_infected_critical = [4.95, (4.95 + 20.58) / 2., 20.58]
    # transmission_prob = [0.02, (0.02 + 0.02) / 2, 0.2]
    # recovered_per_infected_no_symptoms = [0.15, (0.15 / 0.2) / 2, 0.2]
    # risk_of_infection_from_symptomatic = [0.05, (0.05 + 0.2) / 2, 0.2]
    # severe_per_infected_symptoms = [0.006, (0.006 + 0.25) / 2, 0.25]
    # critical_per_severe = [0.05, (0.05 + 0.45) / 2, 0.45]
    # deaths_per_critical = [0.004, (0.004 + 0.7) / 2, 0.7]

    incubation_time = [5., 6.7]
    time_infected = [5.4, 8.4]
    # time_infected = [6]
    # old serial interval
        # 0.5 * 2.67 + 0.5 * 5.2,
        # 0.5 * 4.00 + 0.5 * 5.2,
    serial_interval = [4.2675]
    time_infected_severe = [4., 13.]
    # time_infected_severe = [8.5]
    time_infected_critical = [4.95, 20.58]
    # time_infected_critical = [12.5]
    transmission_prob = [0.02, 0.2]
    recovered_per_infected_no_symptoms = [0.15, 0.2]
    risk_of_infection_from_symptomatic = [0.05, 0.2]
    severe_per_infected_symptoms = [0.006, 0.25]
    # severe_per_infected_symptoms = [0.125]
    critical_per_severe = [0.05, 0.45]
    # critical_per_severe = [0.2]
    deaths_per_critical = [0.004, 0.7]
    # deaths_per_critical = [0.35]

    parameter_combos = list(itertools.product(
        incubation_time,
        time_infected,
        serial_interval,
        time_infected_severe,
        time_infected_critical,
        transmission_prob,
        recovered_per_infected_no_symptoms,
        risk_of_infection_from_symptomatic,
        severe_per_infected_symptoms,
        critical_per_severe,
        deaths_per_critical
    ))

    # Adds mean
    parameter_combos.append(
        [
            np.sum(incubation_time) / 2,
            np.sum(time_infected) / 2,
            np.sum(serial_interval),
            np.sum(time_infected_severe) / 2,
            np.sum(time_infected_critical) / 2,
            np.sum(transmission_prob) / 2,
            np.sum(recovered_per_infected_no_symptoms) / 2,
            np.sum(risk_of_infection_from_symptomatic) / 2,
            np.sum(severe_per_infected_symptoms) / 2,
            np.sum(critical_per_severe) / 2,
            np.sum(deaths_per_critical) / 2
        ]
    )

    death_measurements = []
    names = []
    num_samples = len(parameter_combos)
    # num_samples = 100
    for i in range(num_samples):
        ################################################################################
        # Parameters
        ################################################################################
        for age_group in age_groups:
            model.parameters.IncubationTime[age_group] = parameter_combos[i][0]
            model.parameters.TimeInfectedSymptoms[age_group] = parameter_combos[i][1]
            model.parameters.SerialInterval[age_group] = parameter_combos[i][2]
            model.parameters.TimeInfectedSevere[age_group] = parameter_combos[i][3]
            model.parameters.TimeInfectedCritical[age_group] = parameter_combos[i][4]
            model.parameters.TransmissionProbabilityOnContact[age_group] = parameter_combos[i][5]
            model.parameters.RecoveredPerInfectedNoSymptoms[age_group] = parameter_combos[i][6]
            model.parameters.RiskOfInfectionFromSymptomatic[age_group] = parameter_combos[i][7]
            model.parameters.SeverePerInfectedSymptoms[age_group] = parameter_combos[i][8]
            model.parameters.CriticalPerSevere[age_group] = parameter_combos[i][9]
            model.parameters.DeathsPerCritical[age_group] = parameter_combos[i][10]

        death_measurements.append([
            model.parameters.IncubationTime[A0].value,
            model.parameters.TimeInfectedSymptoms[A0].value,
            model.parameters.SerialInterval[A0].value,
            model.parameters.TimeInfectedSevere[A0].value,
            model.parameters.TimeInfectedCritical[A0].value,
            model.parameters.TransmissionProbabilityOnContact[A0].value,
            model.parameters.RecoveredPerInfectedNoSymptoms[A0].value,
            model.parameters.RiskOfInfectionFromSymptomatic[A0].value,
            model.parameters.SeverePerInfectedSymptoms[A0].value,
            model.parameters.CriticalPerSevere[A0].value,
            model.parameters.DeathsPerCritical[A0].value,
        ])

        baseline_home = np.array(
            [
                [0.4413]
            ]
        )
        contacts = ContactMatrix(np.r_[baseline_home])
        contacts.add_damping(Damping(coeffs=np.r_[0.7], t=30.0, level=0, type=0))

        # print('model.parameters.ContactPatterns.cont_freq_mat')
            # print(model.parameters.ContactPatterns.cont_freq_mat)
            # print('done with model.parameters.ContactPatterns.cont_freq_mat')
        model.parameters.ContactPatterns.cont_freq_mat[0] = contacts
        model.apply_constraints()

        # print('starting sim')
        sim = secir.Simulation(model, t0=0., dt=0.1)
        sim.advance(tmax=1e4)
        # print('finished sim')
        output = secir.interpolate_simulation_result(sim.result, [0, 7, 14, 21, 28])  # .as_ndarray().copy()

        infection_state_count = 10
        vars = [
            "skip Suspectible",
            "skip Exposed",
            "skip Infected asymptomatic",
            "Infected asymptomatic reported",
            "skip Infected symptomatic",
            "Infected symptomatic reported",
            "Hospitalized",
            "ICU",
            "Recovered",
            "Dead"
        ];


        # print('sample',i)
        output = output.as_ndarray().copy()
        names = [
            "o#Total dead (log base 10)",
            f"o#Dead after {output[0][4]} days (log base 10)",
            f"o#Dead after {output[0][3]} days (log base 10)",
            f"o#Dead after {output[0][2]} days (log base 10)",
            f"o#Dead after {output[0][1]} days (log base 10)",
            # f"o#Dead after {output[0][5]} days (log base 10)",
            # f"o#Dead after {output[0][6]} days (log base 10)",
            # f"o#Dead after {output[0][7]} days (log base 10)",
        ]
        death_measurements[-1].append(np.log10(output[-1, -1]))
        death_measurements[-1].append(np.log10(output[-1, 4]))
        death_measurements[-1].append(np.log10(output[-1, 3]))
        death_measurements[-1].append(np.log10(output[-1, 2]))
        death_measurements[-1].append(np.log10(output[-1, 1]))
        # death_measurements[-1].append(np.log10(output[-1, 5]))
        # death_measurements[-1].append(np.log10(output[-1, 6]))
        # death_measurements[-1].append(np.log10(output[-1, 7]))

        # for k in reversed(range(infection_state_count)):
        #     if "skip" in vars[k]:
        #         print("skipping", vars[k])
        #         continue
        #     total_dead = 0
        #     for i in range(num_age_groups):
        #         # names.append("o#" + vars[k] + f"(Age group {i})")
        #         names.append("o#" + vars[k])
        #         death_measurements.append(output[k + infection_state_count * i])
        #
        # deaths.append([
        #               ] + death_measurements)

    columns = [
        # 'IncubationTime',
        # 'TimeInfectedSymptoms',
        # 'SerialInterval',
        # 'TimeInfectedSevere',
        # 'TimeInfectedCritical',
        # 'TransmissionProbabilityOnContact',
        # 'RecoveredPerInfectedNoSymptoms',
        # 'RiskOfInfectionFromSymptomatic',
        # 'SeverePerInfectedSymptoms',
        # 'CriticalPerSevere',
        # 'DeathsPerCritical',
        'IT',
        'TI Symptomatic',
        'Serial Interval',
        'TI Severe',
        'TI Critical',
        'TP',
        'RpI',
        'RI',
        'SpIS',
        'CpS',
        'DpC',
    ]


    deaths = ['i#' + c for c in columns] + names
    deaths_pd = pd.DataFrame(data=death_measurements, columns=deaths)
    deaths_pd = deaths_pd[['i#IT', 'i#TI Symptomatic', 'i#TI Severe', 'i#TI Critical', 'i#TP', 'i#RpI', 'i#RI', 'i#SpIS', 'i#CpS', 'i#DpC'] + names]
    deaths_pd.to_csv('S1_table.csv', index=None)
    deaths_pd_2 = deaths_pd[['i#TP', 'i#DpC']+names]
    deaths_pd_2.to_csv('S2_table.csv', index=None)
```
