## Supplementary material for "MooViE – Engine for single-view visual analysis of multivariate data": run.sh

```
#!/bin/bash

# assumes a locally installed moovie in PATH
# MOOVIE_EXEC="moovie"
# assumes the moovie-desktop docker img is available (e.g. from the moovie-desktop-docker script)
# MOOVIE_EXEC="docker run --rm -ti --user "$(id -u)" -v ${PWD}:/home  moovie-desktop moovie"
# assumes the docker image from the registry
MOOVIE_EXEC="docker run --rm -ti --user "$(id -u)" -v ${PWD}:/home jugit-registry.fz-juelich.de/ibg-1/modsim/moovie/moovie:v1-0-0 moovie"

for i in S2_1*.csv;
do
    $MOOVIE_EXEC -c MOOED.conf -o "${i%.*}.svg" "${i}"
done

for i in S2_2*.csv;
do
    $MOOVIE_EXEC -c RED.conf -o "${i%.*}.svg" "${i}"
done
```
