## Supplementary material for "MooViE – Engine for single-view visual analysis of multivariate data": README.md

# S3: Visual Assessment of Algorithmic Multi-objective Optimizer Performances

To simulate the raw data execute
```
python3 S3_script.py
```

Once the data has been generated (or the existing data is used), the MooViE figures are generable by executing run.sh.
