## Supplementary material for "MooViE – Engine for single-view visual analysis of multivariate data": S3_script.py

```
import pandas
import numpy as np
import matplotlib.pyplot as plt
from pymoo.problems import get_problem
from pymoo.visualization.scatter import Scatter
from pymoo.algorithms.moo.nsga2 import NSGA2
from pymoo.algorithms.moo.moead import MOEAD
from pymoo.util.ref_dirs import get_reference_directions
from pymoo.algorithms.moo.sms import SMSEMOA
from pymoo.termination import get_termination
from pymoo.optimize import minimize

plt.rcParams["axes.labelsize"] = 25

problems = []

dtlz5 = get_problem("dtlz5", n_var=4, elementwise_evaluation=True)
pf = dtlz5.pareto_front()
Scatter(angle=(20, 20), figsize=(15, 10), axis_style={"fontsize": 25}).add(pf).save("pareto_dtlz5.png")
problems.append(("dtlz5", [(0., 1.), (0., 1.), (0., 1.)], dtlz5))

plt.rcParams["axes.labelsize"] = 30
dtlz7 = get_problem("dtlz7", n_var=4, elementwise_evaluation=True)
pf = dtlz7.pareto_front()
Scatter(angle=(20, 20), figsize=(15, 15)).add(pf).save("pareto_dtlz7.png")
problems.append(("dtlz7", [(0., 1.), (0., 1.), (2.6, 6.2)], dtlz7))

termination = get_termination("n_gen", 100)

ref_dirs = get_reference_directions("uniform", 3, n_partitions=20)
for prob_name, ranges, problem in problems:
    print(f"problem {prob_name}")
    for alg_name, algorithm in [("nsga2", NSGA2()),
                                ("moead", MOEAD(ref_dirs,
                                                seed=1)),
                                ("smsemoa", SMSEMOA())]:
        print(f"{alg_name} with population size = {algorithm.pop_size}")
        res = minimize(problem,
                       algorithm,
                       termination,
                       seed=1,
                       save_history=True,
                       verbose=True)

        df = pandas.DataFrame(np.column_stack((res.X[:100],
                                               res.F[:100])),
                              columns=[f"i#x{i + 1}(0, 1)" for i in range(res.X.shape[1])] +
                                      [f"o#f{i + 1}{ranges[i]}" for i in range(res.F.shape[1])])
        df.to_csv(f"{prob_name}_{alg_name}.csv", index=False)
```
