## Supplementary material for "MooViE – Engine for single-view visual analysis of multivariate data": README.md

# generating suplementary figures

The subfolders contain all needed data to generate the supplemenatry figures. For that each subfolder contains as `run.sh` script.

Potentially, the `MOOVIE_EXEC` has to be adjusted to an appropriate location. The script already contains 3 common cases

1. A locally build MooViE
    - `MOOVIE_EXEC="moovie"`
2. A docker image generated from the `moovie-desktop-docker` script
    - `MOOVIE_EXEC="docker run --rm -ti --user "$(id -u)" -v ${PWD}:/home  moovie-desktop moovie"`
3. DEFAULT: The official docker image from the registry. As this pins the version, it guarantess maximal reproducibility.
    - `MOOVIE_EXEC="docker run --rm -ti --user "$(id -u)" -v ${PWD}:/home jugit-registry.fz-juelich.de/ibg-1/modsim/moovie/moovie:v1-0-0 moovie"`

Uncomment or add, as you need.

For case 1 (Compartmental Models in Epidemiology) and case 3 (Visual Assessment of Algorithmic Multi-objective Optimizer Performances) additional scripts are 
present to reproduce the raw data. For details please refer to the respective `README.md` files. Case 2 (Experimental Design for Isotope Tracer 
Experiments) relies on previously published data and data generation is described in the corresponding publications.

To generate the images for a specific case run

- `run.sh`
- `for i in *.svg; inkscape --export-type="png" -d 200 $i; done`
- `rm *.svg`
- `for i in *.png; do convert $i -trim $i; done`
